# Computational Investigation Of Structural Interfaces Of Protein Complexes With Short Linear Motifs

**DOI:** 10.1101/2020.03.27.012864

**Authors:** Raghavender Surya Upadhyayula

## Abstract

Protein complexes with short linear motifs (SLiMs) are known to play important regulatory functions in eukaryotes. In this investigation, I have studied the structures deposited in PDB with SLiMs. The structures Were grouped into three broad categories of protein-protein, protein-peptide and the rest as others. Protein-peptide complexes Were found to be most highly represented. The interfaces Were evaluated for geometric features and conformational variables. It was observed that protein-protein and protein-peptide complexes show characteristic differences in residue pairings, which Were quantified by evaluating normalized contact residue pairing frequencies. Interface residues adopt characteristic canonical residue conformations in the Ramachandran space, with a pronounced preference for positive ϕ conformations. It was observed that phosphorylated residues have an unusual propensity to adopt the unusual positive ϕ conformations at the interface.

## Introduction

The interaction between a protein and a partner in the cell is quite complicated, the rules for which are being revealed in layers with experiments, computational calculations and predictions. This type of interaction involves binding of the polypeptide chains through an interface formed between both the chains. The interface between the chains is stabilized by interactions such as hydrogen bonds, van der Waals forces, salt bridges and ionic interactions. The study of these interactions is to identify binding sites and predict which interactions are energetically favorable and help in understanding the construction of binding site at chemical and atomic level. The growing number of functional protein complex structures in the Protein Data Bank (PDB) has spurned the interest of many computational investigators to decipher the molecular determinants of complex formation.^1,2^

The protein-protein interaction data obtained from experiments is prone to contain false positives / false negatives. This adds an unknown amount of bias in predictions, if the data Were to be used *in toto* without any manual intervention.^3^ Computational methods can be exploited for understanding the various interaction networks derived from experimental methods. In pioneering studies by the Barabasi and coworkers, network topology was found to be responsible for robustness. In this picture some proteins serve as ‘hubs’ with dense connections and the rest are at the edges with fewer connections.^4–6^ Extrapolation of computational network models generated to PPIs of the whole proteome of an organism requires understanding unique PPIs and the functional significance with the observations *in vitro or in vivo*. Recently, Barabasi group have proposed a novel approach of identifying viable PPIs by predicting interactions with proteins which share similar partners using network paths of length 3 (L3).^7^ The study of interfaces is rewarding in not only understanding the environment of protein binding sites but also provides a handle on engineering new interactions.^8–15^ These interfaces have been studied at both protein and domain levels and a large number of tools, databases and interface datasets are available for analyzing PPIs using computational methods.^16–19^ Many visualization tools are being actively developed and are discussed in many articles. More than 100 databases are available for PPIs and are growing by every year.^20^

The functional interaction modules in the intrinsically disordered regions (IDRs) of higher eukaryotic proteomes are rich in regulatory interaction interfaces. Lack of bulky hydrophobic residues in IDRs leads to a non-classical structure-function paradigm in these sequences in complete contrast to globular, structured proteins.^21,22^ The current view of a proteome is that of a module comprising of proteins with structured and disordered regions. Structured proteins with disordered regions provide the functional variety and diversity regulating the cellular processes.^23^ Disordered regions are good targets for post translational modifications (PTMs) which present a multitude of functional states of such proteins in the cell.^24,25^ As much as 44% of protein coding genes contain disordered segments in human proteome with sequences comprising of more than 30 residues.^26^ A major class of disordered segments which are involved in interaction interfaces are the compact and degenerate modules known as short linear motifs (SLiMs) comprising of 3-10 residues.^27^ Extracting or predicting these motifs, either experimentally or computationally, is challenging because of their weak binding and high degree of degeneracy. These disordered regions are classified to be involved in three major categories of functional importance of transient binding (chaperones, PTMs), permanent binding (effectors, assemblers, scavengers) and no binding (entropic chains).^28^ Dunker and coworkers have arrived at 28 unique functions of the IDRs based on literature analysis of 150 proteins containing IDRs.^29^ ELM and MiniMotif are two approaches to annotate linear motifs found in IDRs and their binding partners.^30^

Rapid increase of available protein structures provides an excellent opportunity to study SLiMs directly from their 3D structures. SLiMs are considered as a functional module within IDRs, and with the support from the flexible adjoining disordered regions are known to bind the surfaces of globular domains compactly.^18,31,32^ The short nature of SLiMs makes them amenable to a high propensity for evolving convergently and emerge in unrelated proteins.^33,34^ In one of the initial attempts to study and predict SLiMs on protein structures, Hugo et al., have discussed the importance of SLiM detection on protein-protein interfaces.^35^ Their approach was to find *de novo* SLiMs on protein interfaces, which involved domain interface clustering and extraction of putative SLiMs. The *average* protein-protein interface *area* defined in their study was around 800 Å^2^ or greater, agreeing with the accepted limit for domain interfaces.^36^ Our understanding of the fundamental principles of structural association of SLiM mediated interactions suffer from the difficulties associated with the experimental identification and validation of SLiMs. An important advance in the field was the identification of molecular recognition features (MoRFs) within IDRs. Uversky and colleagues studied loosely structured regions (10-70 residues) within IDRs which undergo disorder-to-order transition upon binding with their partner. They came upon classifying MoRFs into three fundamental categories of α-MoRF, β-MoRF and ι-MoRFs depending upon the structure adopted upon binding (α-helix, β-strands and irregular).^37^ Another important contribution was from Luis Serrano to decipher that the intrinsic propensity of ϕ torsion of each amino acid shows significant difference in specific secondary structures.^38^ It was established conclusively that the side chain is responsible for the observed relative ϕ preferences. In fact, secondary structure propensities for specific types of structurally solved regions of polypeptide chains have led to pioneering insights into the intrinsic preferences of individual residues.^39,40^ Motivated by these studies, I posed the following question whether there is some critical information in ELM database which can aid in the design of targeted inhibitors by computational and statistical means. With the growing number of structures deposited in PDB, the need to study and analyze the SLiM interfaces is scientifically appealing.

Specifically, in this report I focus on analyzing the characteristics of structural interfaces of complexes in the eukaryotic linear motif (ELM) database and evaluate the intrinsic preferences of the interface residues. Residue level conformational features at complex interfaces and the role of weak interactions in shaping the interfaces are investigated. These are contrasted with the trends seen in the known classes of protein-complexes. In addition, I observe unusual conformations of canonical residues and phosphorylated variants of Ser/Thr/Tyr (pSer/pThr/pTyr) at structural interfaces and evaluated their intrinsic conformational propensities. To my understanding, for the first time the propensities of positive ϕ conformations are reported for SLiMs. I have compared and contrasted the calculated propensities with models and datasets reported in the literature for residues at interfaces. I believe that this knowledge of all possible geometric and residue specific dependencies should aid in design of specific inhibitors and also help in the development/refinement of prediction algorithms.

## Methods

### Dataset Creation

A list of complex structures from PDB Were procured from the ELM database (http://elm.eu.org). The initial list comprised of 452 complex structures solved by X-ray crystallography (X-ray) and Nuclear Magnetic Resonance (NMR). I chose to work with representative structures at 90 % sequence identity. This resulted in 231 structures solved by X-ray crystallography with resolution better than 3.0 Å.

### Evaluation of dihedrals and interface interactions

The conformation of the residues {*ϕ*, *ψ*, *χ*^1^, *χ*^2^, *χ*^3^ *χ*^4^} were calculated using the program *dangle* from the Richardson lab. Weak interactions at the interfaces were curated by obtaining the information from PDBSum. Independently, I calculated the protein interactions by manually viewing and cleaning the structures using Maestro v11.7.012. Briefly, I load the structures in Maestro interface and prepare the structure using the Protein Preparation Wizard using default options. Hydrogen bond assignment includes changes in protonation states of Asp, Glu and tautomeric states of His residues (HID/HIE) to maximize H-bond network. In case of NMR structures, all the models Were used for calculations as they would harbor subtle energetically stable conformational variations between models. Protein Interaction Analysis module was used to calculate interactions. Polypeptide chains Were split into protein and peptide interacting groups by visualizing them in the software. The parameters used in the calculation of interaction parameters include finding neighbors within 4 Å of interface residues. Hydrogen bonds are found based on the following definitions: D-H…A-X is the hydrogen bond entity with D standing for donor, A for acceptor and X is a heavy atom. The minimum acceptor angle is 90°, minimum donor angle is 120° and the maximum distance between heavy atoms is 3.5 Å. Aromatic-aromatic interactions are quantified by measuring the centroid to centroid distance up to a maximum of 6.5 Å. The van der Waals clashes are listed based on the allowable overlap of atoms (R_A_ + R_B_ − R_AB_) at 0.4 Å. Further data processing was done in R and Python.

### Evaluation of Contact Residue Pair Preferences

I have followed the refined and optimized methodology proposed by Blundell and coworkers for calculating contact pairing preferences.^41^ The frequency of pairwise residue interactions (*P*_*ij*_) Were derived by using the equation:

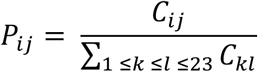

where *C*_*ij*_ represents the number of times residue type *i* is found engaging with residue type *j*. I then calculated individual frequencies reflecting the residue composition *and normalized taking into account buried SASA*:

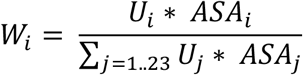

here *U*_*i*_ represents number of residues engaged in contacts. I then calculated log odds ratio of the *observed* frequency to the *expected* frequency as:

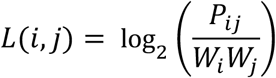

This is a commonly used measure to quantify interaction preference at interfaces, with the understanding that larger residues have greater surface area making them better candidates for interaction with one another.^42^ With residue sizes are accounted for in the above definition, the bias in the calculated interaction preference is lifted.

## Results and Discussion

### Composition of Dataset

Unique polypeptide chains were identified from the sequence of each polypeptide chain with coordinates in the structure file. Polypeptide chains with less than or equal to 30 residues were treated as peptides in this analysis. The reason for using such a large value for peptide length definition is arbitrary and to an extent is based on our observation of synthetic peptides of length > 30 in both PDB and Cambridge Structural Database^43^. In addition, SLiMs are known to be short (3-10 amino acids) and >30 residues are treated as IDRs.

The structures were divided into three classes as protein-protein, protein-peptide and others. The third category of 20 structures consists of mostly monomeric and small molecule bound proteins, with one example of a peptide-peptide complex (2JO9) and one multiprotein-peptide complex (3VI4). These structures are excluded from analyses, as our focus is on understanding the residue preferences and conformations of dimeric complexes. I observe that majority of the structures, 76%, belong to protein-peptide complexes and solved by X-ray diffraction (Figure 1(a)). Protein-protein comprises of 15% of the dataset. I choose to impose a flexible B-factor based filter to retain structures for further analyses. Specifically, I choose to keep structures with an average B factor for the entire structure to be ≤60.0 Å^2^. Contrary to this, the analyses focused on deriving accurate parameters for modeling rely on extremely stringent B-factor upper limit to account for the precision of atoms in the residues. Relaxing the condition on structure B-factor allowed us to account for the flexible regions of the structure which are generally discarded in geometric analyses. This is from the viewpoint that the information contained in the coordinates, even though imprecise can provide valuable insights in the light of studying intrinsically disordered regions of proteins. I have plotted R_free_ of the structures on a dual axis (Figure 1(b)). Broadly, there are two clusters with R_free_ in the range of (0.18, 0.22) to (0.24, 0.30). The composition of the interfaces is described by pairwise interactions of individual residues in three categories. I evaluated the interactions at the interface based on the parameters of hydrogen bonds, van der Waals clashes, disulfides, salt bridges and π-π stacking (Figure 1(c)). Domain composition was obtained by mapping SCOP domains and regions onto the structures (Figure 1(d)).

**Figure 1.**
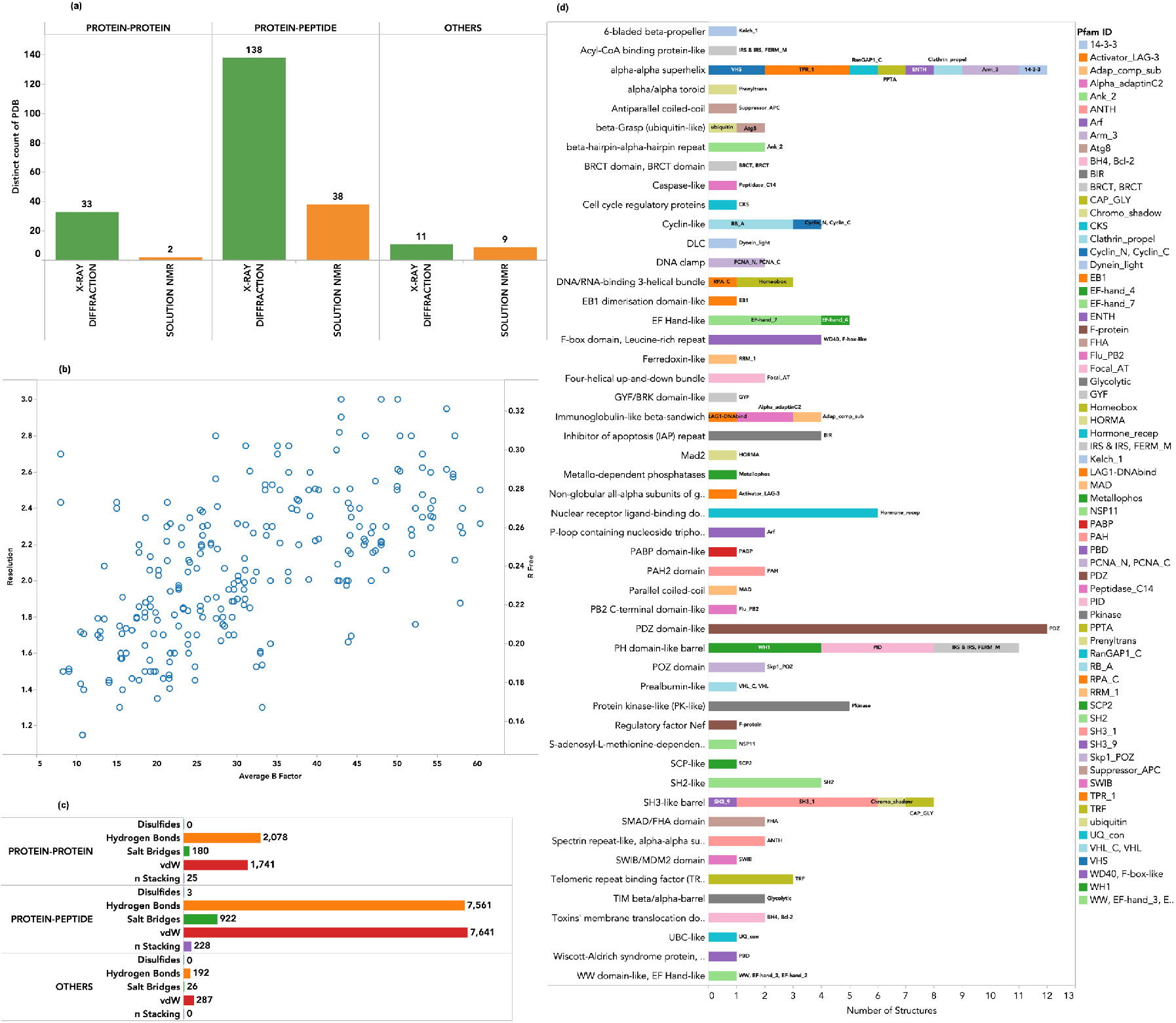
Counts of (a) complex structures by types and methods used to solve them; (b) Resolution and R-free versus Average B-factor of structures, (c) Counts of various interactions in three categories of complexes and (d) SCOP-IDs and Pfam IDs observed in the structures.

Of those structures with reliable mapping of SCOP domains, α-α superhelix, PDZ domain-like and PH-domain-like barrel are the most frequent. These are followed by SH3-like barrels, protein-kinase like, cyclin-like and EF-hand like. The coverage of unique domains, although quite limited, is significant because of the regulatory roles which are implied for these important domains.

### Interface Interactions

I observe that hydrogen bonds and van der Waals interactions are the major forces cementing the polypeptide chains at the interfaces in all the three categories. Salt bridges are important in protein-protein and protein-peptide interfaces.^44^ This is followed by stacking of aromatic rings in π…π interactions. This is in agreement with previous observations that in PPI non-polar and aromatic side chain containing residues are the major players at the protein interfaces.^15,45^ The differences in the interface composition and preferences of obligate and non-obligate complexes have been discussed in earlier studies showing that the differences in the strength of interactions is essentially due to the nature of residues.^46^ The former relies on forming relatively more hydrophobic contacts and the latter on salt bridges and hydrogen bonds.^1,47^ In our dataset I observe that hydrogen bonds make up 51.6% and 46.2% in protein-protein and protein-peptide interfaces. Salt bridges share 4.4% and 5.6% and π…π interactions make up 0.6% and 1.3%, respectively. These differences tend to indicate that the trend is the same in both the complex data sets.

### Buried Solvent Accessible Surface Area (SASA)

Buried SASA is an excellent geometric measure of the size of the protein interfaces. It has been established that protein-protein interfaces are typically in the range of 1500 Å^2^ and relatively flat with pockets, crevices and indentations.^47,48^ Peptides tend to be shorter and accordingly bury surfaces of the order of 500 Å^2^. The fraction of buried SASA of a residue that is buried by the interaction with other residues of the partner provides clues to the further classification of interface residues into core and rim categories (Figure 2). I categorized the interfacial residues with buried SASA fraction ≤30% as ‘rim’ and the rest as ‘core’. The pie section corresponds to the sum of buried SASA for that particular residue in observed instances. I see that the majority of the interface residues are in the core with the residues PTR (pThr), TPO (pTyr) and SEP (pSer) observed at rim as well. Amongst canonical residues, Lys, Asp, Gln, Glu, have explored the rim regions at the interfaces. The longer and charged sidechains of these residues appear to explore the interface region flexibly.

**Figure 2.**
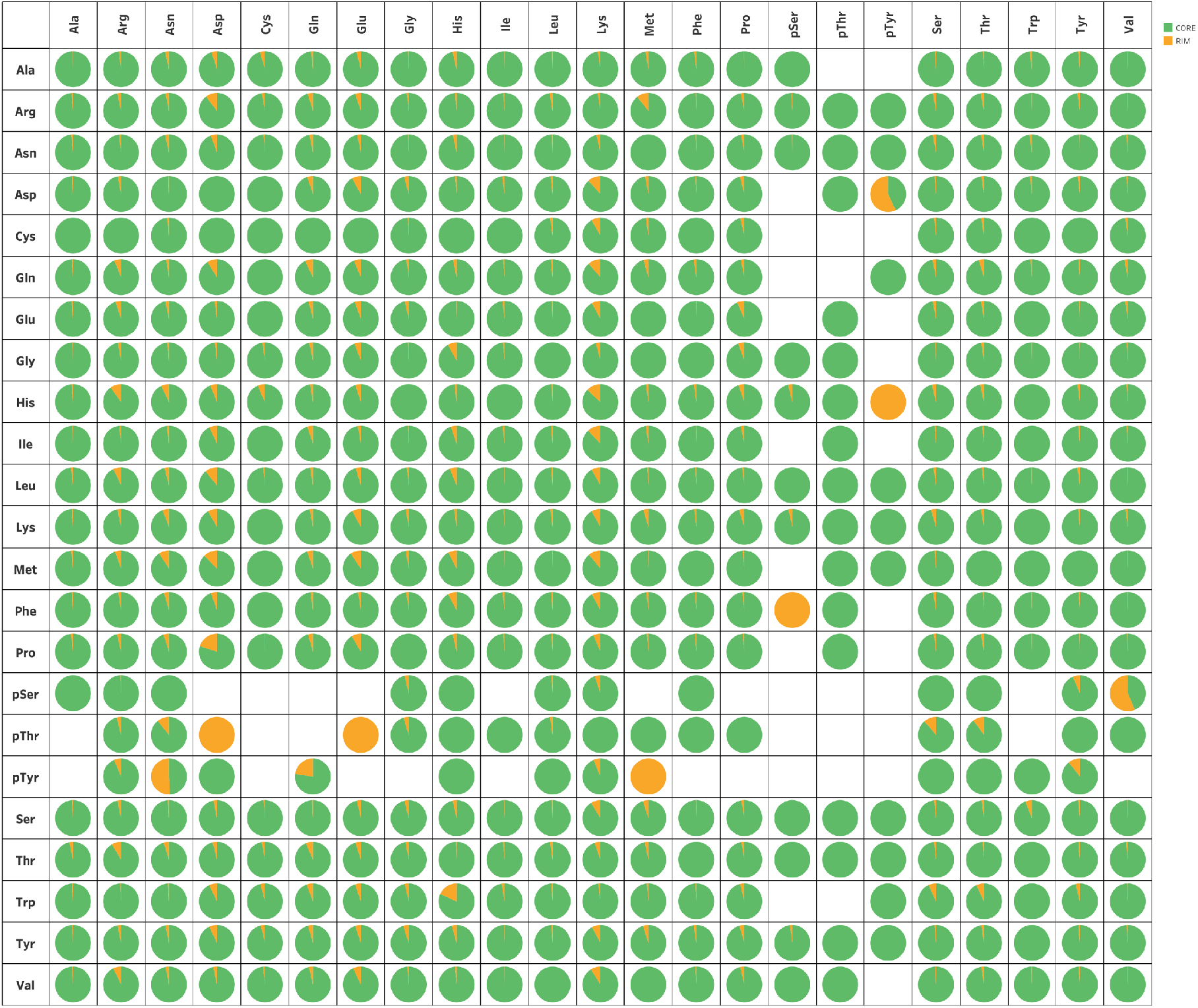
Buried solvent accessible surface area (SASA). Residues on chain 1 are on *x*-axis and residues in chain 2 on *y*-axis

### Conformational analysis of residues at the interfaces

The conformations of residues at the interface of the complexes provides an insight into the intrinsic preference of individual residues to adopt specific conformations. I subdivide the backbone conformations (ϕ, ψ) of individual residues into two categories as discussed above (Figure 3). Ramachandran maps of residues grouped based on their chemical nature are also provided (Supplementary Figure 1).

**Figure 3.**
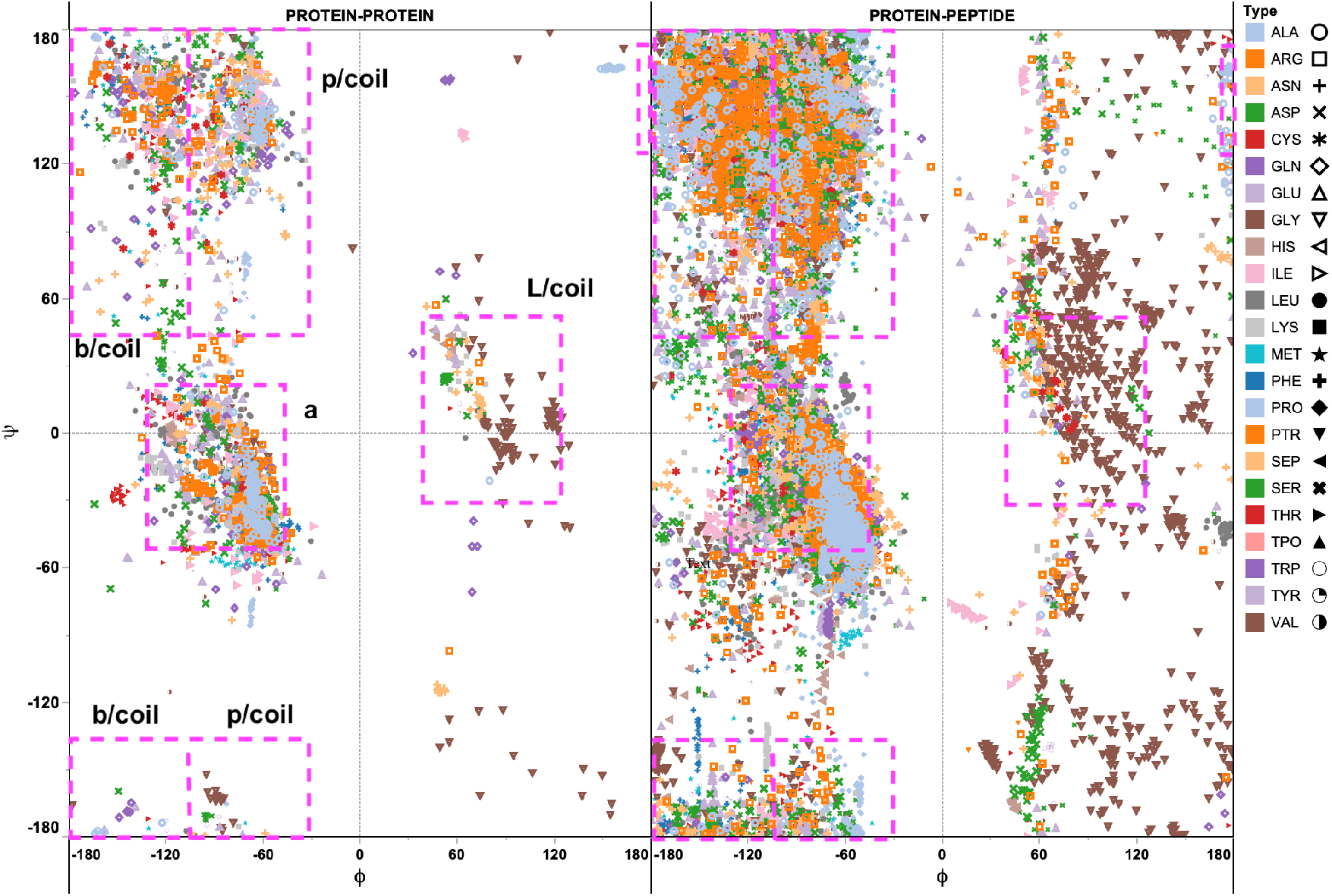
Ramachandran maps of interface residues in ELM dataset. Boxed regions (magenta) correspond to coil regions described in Swindell’s work.

The conformations observed are consistent with previously published preferences for canonical residues at protein-protein interfaces.^49^ The backbone torsions appear to follow similar trends in both the categories. An important observation is that most of the canonical residues adopt unusual positive ϕ conformations. The intrinsic preferences calculated by Swindell’s et. al., for coil regions serves as a reference for understanding these observations.^40^

### Positive ϕ Conformations of Residues at the Interfaces and their propensities

Apart from Gly which is well known to adopt positive ϕ conformations, I note that phosphorylated modifications of threonine and tyrosine have a high propensity to adopt unusual conformations at the interface positions. Ala, Arg, Asn, Cys, Leu, Ser, Thr, and Trp also show propensity to be at interface with positive ϕ conformation in protein-peptide than in protein-protein complexes. His, Ile and Lys in protein-protein complexes show a similar trend. The rest have non-discernible differences for comparison (Supplementary Figure 2).

### Hydrophobics

In the case of hydrophobic residues in the protein-protein complex 2LNH solved by NMR I observe Ala-162 and Ile-159 on chain C to adopt positive ϕ values. This is a hetero-trimer of three different polypeptide chains and chain C represents secreted effector protein EspF(U) from *E.coli*. A GTPase domain binding ligand is located in this protein with ELM instance ELMI003350. ELM is located to the N-terminus of the two residues on a helix. The residues exist on a long loop interfacing with the β-hairpin loop of chain B. In 2O8G, the Leu-289 at the C-terminus of the protein. In 2W96 Trp-150 on chain and in 3L4F, Ala-640 on chain A is observed to adopt a positive ϕ. The ELM motif (ELMI001573) sequence is adjacent to this residue is a PDZ ligand.

### Charged and Polar

There is a single instance in where Arg adopts positive ϕ. It is part of a protein segment which is having a WW domain of Group 1 and interacts with PPxY motif, where x represents any amino acid. Surprisingly, this ELM motif (LIG_WW_1) has only two reported structures from solved by both X-ray (1EG4) and NMR (2LB1). In both the cases, the Arg residues are in a loop interfacing with the motif containing peptide. In the NMR structure Arg-304 interacts with Tyr-227 and in X-ray Arg-190 interfaces with Arg-7 in the peptide. I rationalize that irrespective of the residue being buried or exposed in a protein-peptide interaction the partner appears to induce the unfavorable conformation in the protein residues.

### Positive ϕ in pTyr and pThr containing interfaces

It is interesting to note that 14% pTyr residues in our data adopt positive ϕ in the non-L/coil region. These are seen in the NMR structure of FHA1 domain of Rad53 complexed with a biologically relevant phosphopeptide derived from Madt1 (2A0T, Figure 4a). I believe that the unusual conformation is induced in pTyr-169 to establish a stable side-chain hydrogen bond with the interface residues Asn-86 and a backbone hydrogen bond with Ser-85. This correlates with the observation of Serrano that backbone conformations can influence the structural interactions of interacting residues.^38^ A similar proportion of pThr residues (17%) are seen to adopt positive ϕ values. In this case, the values are spread to a greater extent in the positive ϕ quadrants of Ramachandran plot (Figure 3). These examples are of proteins/domains involved in regulatory events with immunologically important domains SH2 (2EU0 and 1MW4) and MAPK signaling pathway (2L1C).^50–52^ I observe that pThr with a bulky benzyl ring finds additional stability by interacting with positively charged residues (Lys, Arg) and also by stabilizing ring interaction with His (Figure 4b). Specifically, pThr-759 in 2L1C is seen to fit nicely in the canonical PTB site (Figure 4c). The residues FTNITpY^759^ form the classical consensus ϕxNPx**pY**Shc PTB recognition motif. The stabilization is due to salt bridge formation between pThr-759 and the positively charged residues in the pocket.^53^ Although the structure was used to understand the binding mechanism, I observe that the key to this association lies in the preference of pThr to adopt positive ϕ conformation.

**Figure 4.**
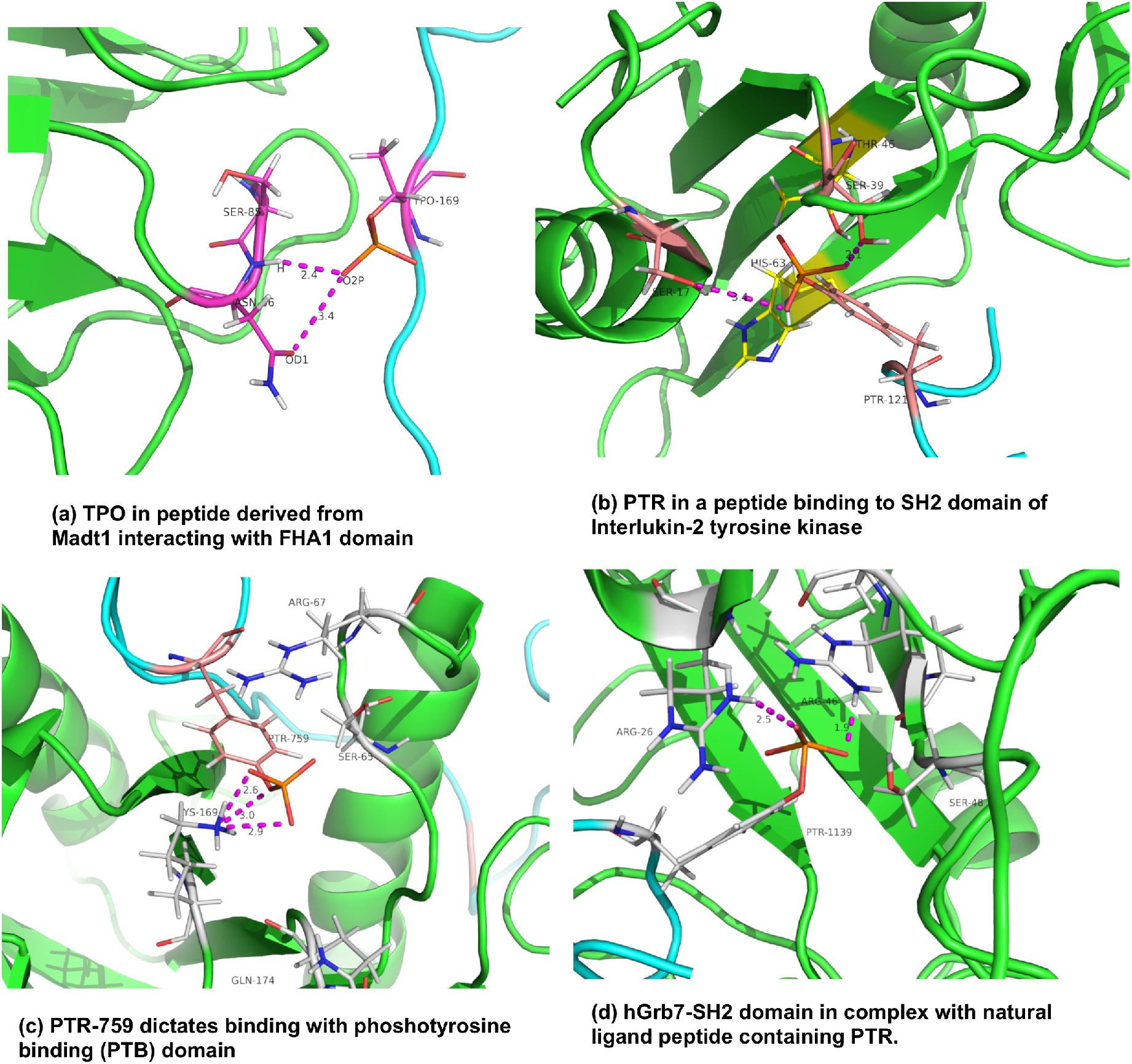
Structural interactions and stability of phosphorylated peptide ligands adopting positive ϕ values. PDB codes for (a) 2A0T, (b) 2EU0, (c) 2L1C and (d) 1MW4

### Propensity of a residue to occur at interface with positive ϕ

Pair potential matrices were proposed for studying the pairing potential of residues across structural interfaces by Sternberg and coworkers. Residue level pair potentials provide a way of a statistical measure of likelihood of that pair occurring and the total likelihood of the structure is the sum of all the individual potentials. As suggested by Sternberg this method shares equivalence with Boltzmann’s law and thereby is related to an estimate of relative free energies for different residue pairings.^42^ The propensity of residue-residue pair contacts for the two groups is calculated and shown (Supplementary Figure 2).

I observe that the natural chemical likeness is preserved with like groups preferring partnership (hydrophobic-hydrophobic, charged-charged etc.). This can be contrasted with those reported in PICCOLO database and that by Glaser wherein Arg-Trp is the most favored pair in protein-protein complexes.^54^ One of the first reports to comprehensively distinguish different types of interfaces into six categories by Burkhard Rost, clearly highlighted the residue-residue pairing preferences to be remarkably different in each class. The residue pair contact definition adopted in this report is similar to that of Blundell and coworkers.^41^ As pointed out by Rost, datasets with smaller data sets is that significance tests become problematic.

All the canonical residues were observed to adopt positive ϕ values, suggesting that the propensity for each residue to adopt such unusual conformations could shed insights into their interaction preference at structural interfaces. The propensities for residues adopting ϕ > 0° were evaluated and correlated with propensity scales devised earlier for loop residues and protein-protein interface residues.^40,41^ A propensity score was devised with our dataset to quantify the observed propensity of interface residues to adopt positive ϕ conformations as below:

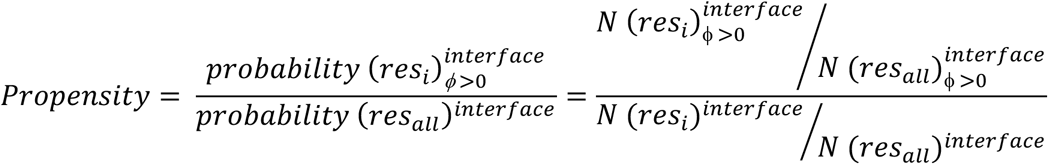

A summary of the *z*-score normalized propensities of each observed interface residue across the five categories comprising of coil (L/coil, b/coil, p/coil) and protein-protein interfaces is shown in Figure 5.

**Figure 5.**
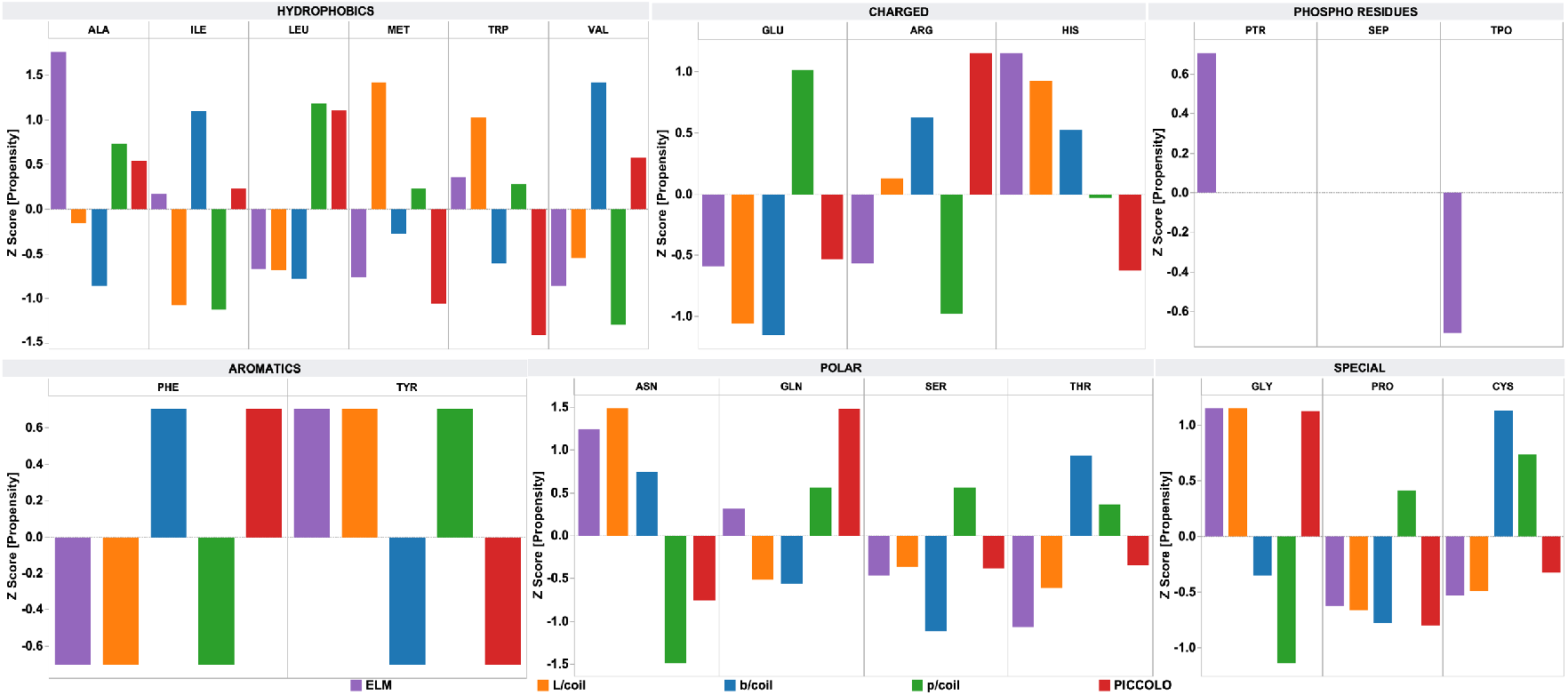
Positive *ϕ* propensity of interface residues in ELM database compared with those found in coil regions (L/coil, p/coil and b/coil) and with those found in interfaces of protein-protein complexes (PICCOLO).

It can be seen that for Ala, Ile, Met, Arg and Gln the *z*-score value of propensities is against that of L/coil data and Trp, Val, Arg, His, Phe, Tyr and Asn have opposite scores to PICCOLO data points. I next attempted to check for correlations with propensities of residues in coil conformations and that of protein-protein interfaces described in monomeric (coil) and protein-protein complexes (PICCOLO) with that of residues adopting positive ϕ conformations.^41^ L/coil region is restricted to positive ϕ regions, with p/coil region having a narrow region in the upper right quadrant of Ramachandran map (Figure 3). Propensity values for canonical residues in our dataset correlate weakly with that of coil regions and those found for protein-protein interfaces. Polynomial equations were seen to fit data points better than linear, logarithmic, exponential, or power regression models (Figure 6). A polynomial of degree 5 is seen to lead to a model which fits well the data points of ELM and L- and p/coil (Supplementary Tables 3-4). It was as expected, since only the L/coil and p/coil regions harbor residues with ϕ > 0°. It is interesting to note that positive ϕ in coil regions of monomeric proteins correlate well with that of positive ϕ conformation of residues in protein-protein and protein-peptide complexes.

**Figure 6.**
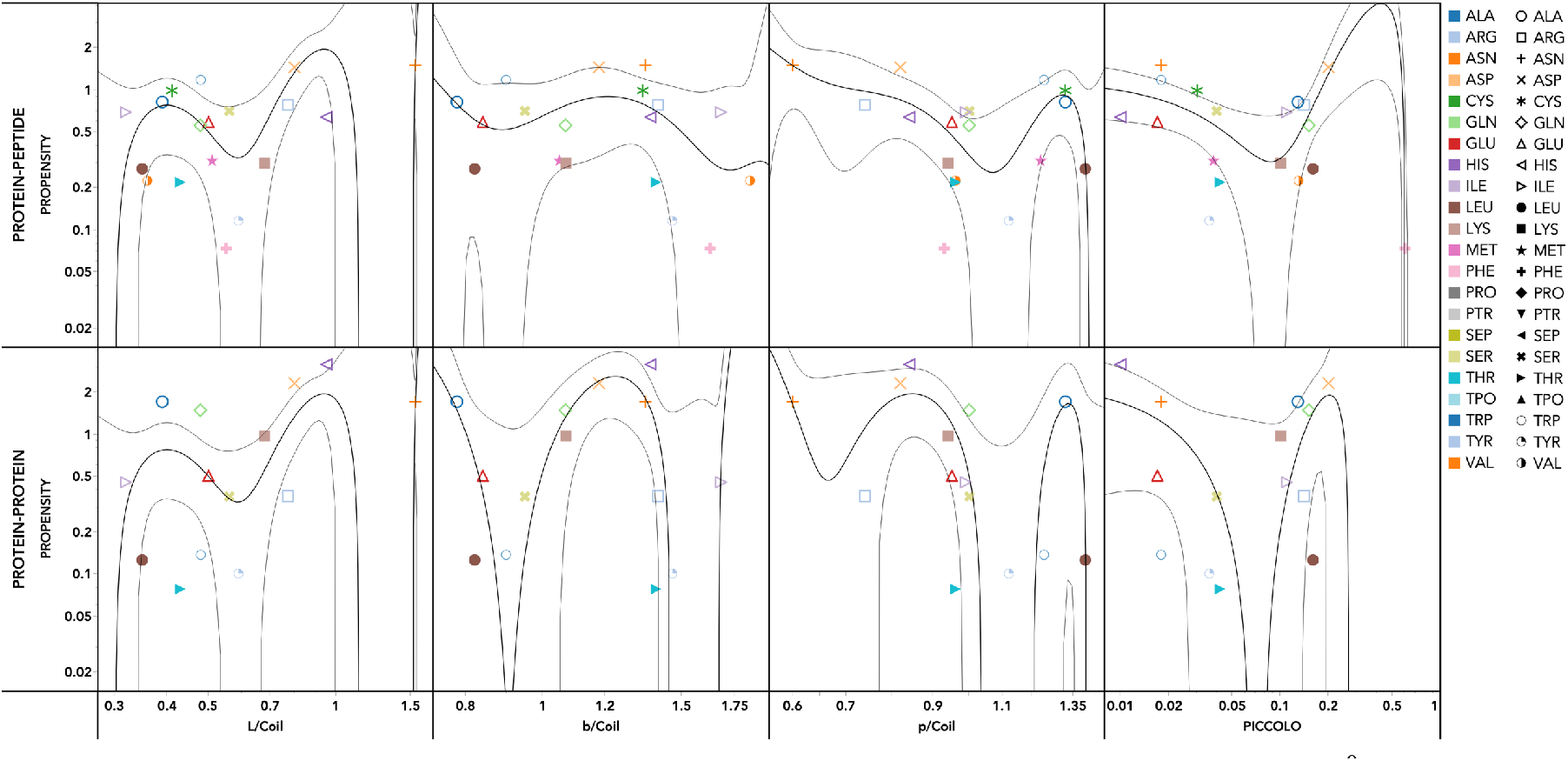
Regression of propensities (log scale) observed for residues adopting ϕ > 0^0^ with those of coil regions and protein-protein interfaces (PICCOLO). 95% CI also drawn as thin magenta black lines. Gly omitted.

## Conclusion

In this investigation I have shown the molecular and structural aspects of distinction between protein-protein and protein-peptide complexes specifically observed in Eukaryotic Linear Motif (ELM) database. The database is manually curated and is repository to many of the well-known functional and regulatory motifs in protein complexes. I evaluated the geometric features of protein interfaces and compared with the known interface descriptors in protein interfaces in complexes and monomeric proteins. Specifically, I observe that at the interface many of the residues have a tendency to adopt ϕ > 0° backbone conformation. I have rationalized these observations by showing that the trend in the conformations of canonical residues adopting ϕ > 0° are in good agreement with the intrinsic (ϕ, ψ) preferences observed for coil residues in monomeric proteins. This result is thought provoking and has implications in designing mimics which can replicate these conformations at the interface, leading to better inhibitors of PPIs.

## Acknowledgements

RSU thank Science and Engineering Research Board (SERB) for awarding Young Scientist Scheme Grant (File No: YSS/2015/001077). Prof. Andy Karplus (Oregon State University, Corvallis, USA) for encouraging RSU to look at unusual positive conformations in protein structures. I thank Dr. M. Krishna Mohan for constant encouragement and support. I thank Birla Institute of Scientific Research (BISR) for providing infrastructural support.

## Additional Information

The authors declare no competing interests related to this work.

## Author Contributions

RSU conceptualized, implemented and analyzed all the aspects of the investigation. RSU wrote the manuscript and prepared all the figures and tables.

